# Molecular Mechanisms for the Biological Storage of Renewable Energy

**DOI:** 10.1101/028381

**Authors:** Buz Barstow

## Abstract

Recent and ongoing discoveries in the field of extracellular electron transport offer the potential to electrically power highly flexible, carbon-fixing microbial metabolisms and generate a rich variety of chemicals and fuels. This process, electrosynthesis, creates the opportunity to use biology for the low cost storage of renewable electricity and the synthesis of fuels that produce no net carbon dioxide. This article highlights recent discoveries on the molecular machinery underpinning electrosynthesis and reviews recent work on the energy conversion efficiency of photosynthesis to begin to establish a framework to quantify the overall energy storage and conversion efficiency of electrosynthesis.

## 1. Introduction

The storage and retrieval of extremely large amounts of renewable energy^1^ and the large scale synthesis of non-carbon-polluting portable fuels^2^ are likely to be key aspects of a future sustainable energy infrastructure. Biological photosynthesis provides an almost complete first draft solution to both of these problems.

Photosynthetic organisms have the remarkable ability to capture sunlight and atmospheric CO_2_ and store both in a variety of energy-dense compounds including starch, sucrose and cellulose using catalysts that self-assemble from abundant elements and function at room temperature and pressure^3^. In addition, biology offers an amazing collection of enzymes that are able to work in concert to transform this fixed carbon to a variety of energy-dense hydrocarbon fuels and fuel precursors including ethanol^4^, butanol^5^, biodiesel^6,7^, branched chain alcohols^8^, medium chain fatty acids^9^ and alkanes^10,11^ that in principle will produce no net CO_2_ over their lifetime^2,12,13^. These enzymatic catalysts can either be incorporated into bacteria and yeast that convert plant biomass into fuels^7,14^, encoded into photosynthetic microorganisms such as algae and cyanobacteria to allow direct conversion of the primary products of carbon fixation into fuels^15^, or perhaps even engineered into plants^16^.

Photosynthesis demonstrates that renewable energy can not only be captured by inexpensive self-assembling catalysts, but also that this can be done at an enormous scale. It is estimated that over the course of a year and across the entire surface of the Earth photosynthesis stores ≈ 3000 exajoules (EJ) (3 χ 10_21_ J) of solar energy as biomass, including crops; forests; algae and subsurface biomass. This corresponds to an average instantaneous energy storage rate of ≈ 100 terawatts (TW)^17^. By contrast, world energy consumption in 2013 stood at only 604 EJ per year (an instantaneous rate of 19 TW)^18^.

Even with the growth in world energy use, the amount of energy channeled by photosynthesis will remain larger for some time to come. Total world energy use is projected to increase to ≈ 865 EJ per year (27.4 TW) by 2040^18^. Should current average rates of increase continue until 2100, world energy consumption could stand at ≈ 1370 EJ per year^18^. Only if we imagine that each of the between 6.9^19^ and 10.9 billion^20^ people who are likely to be alive in 2100 uses energy at the rate that the average American does today (≈ 10 kW)^21^ should world annual energy usage approach or exceed the 3000 EJ channeled by natural photosynthesis per year.

Fortunately, the world is replete with renewable energy. Even when accounting for large increases, world energy consumption rates will constitute only a small fraction of the ≈ 80,000 TW of solar power available at the Earth’s surface^22^. Sourcing a substantial fraction of world energy use from the Sun is a tantalizing possibility. Several classes of technologies offer the possibility of capturing and then storing solar power.

Solar to electrical power conversion technologies are the most advanced. The costs of solar photovoltaics decline at a Moore’s Law pace (it is called Swanson’s Law^23^), while electrical conversion efficiencies continue to rise. The highest recorded light to electrical energy conversion efficiencies currently stand at ≈ 26% for single junction crystalline silicon cell modules, the most widely used material^24^ and ≈ 17% for single-junction CdTe cell modules, the most widely used thin film material^24^. For multi-junction cells under concentrated light, a conversion efficiency record of ≈ 46% is held by a GaInP/GaAs; GaInAsP/ GaInAs cell^24^.

Challenges and opportunities lie in the development of both biological and non-biological technologies to both store this power for later use on the grid or to convert it to energy dense transportation fuels. There has been considerable recent progress in the development of electrochemical catalysts for the conversion of solar electricity to H_2_^25^ and the direct fixation of CO_2_ to CO and formate for use in fuel synthesis^26,27^.

It can be argued that biofuels are the most successful solar power capture and storage method to date. The energy content of all biofuels produced in the US in 2014 stood at 2.25 EJ^28^. By contrast, the integrated solar electrical power delivered in 2014 in the US amounted to only 0.45 EJ^28^. However, substantial challenges lie in the way of increased use. Under field conditions, photosynthesis is only estimated to be able to capture between 0.25^29^ and 1%^30^ of the incident solar flux. This inefficiency has provoked considerable concern over significant further expansion of biofuel production due to the displacement of agricultural production and wilderness and the carbon release incurred by clearing land for energy crops^31,32^. These concerns have encouraged research efforts to improve overall crop yields spanning from traditional crop breeding^33^ to advanced genetic engineering to improve carbon fixation^34^ and biomass characteristics for biofuels^35^.

Two recent trends have encouraged attempts to develop new types of hybrid photosynthesis that combine the metabolic flexibility of biology with the energy capture efficiency of renewable electricity sources, particularly solar electricity. Firstly, a growing appreciation of the physical mechanisms underlying photosynthesis suggests a fundamental mismatch between the ability of the photosynthetic cell to absorb light and its ability to fix carbon. In the wild, photosynthetic organisms appear to exploit this mismatch to dissipate excess light, starving their neighbors of sunlight. While this strategy is advantageous from the viewpoint of evolutionary competition, it works counter to the goal of maximizing solar energy storage. However, while the overall conversion of solar energy to biomass has a low efficiency, this work suggests that far from all of photosynthetic metabolism is inefficient: much of the inefficiency derives from those parts that directly interact with sunlight. Secondly, recent developments in the understanding of extracellular electron transport and autotrophic metabolisms not dependent upon sunlight have begun to build a toolkit of molecular machinery for synthetic biology that will allow the connection of biological metabolism with renewable electricity sources.

Together, these developments suggest the possibility of replacing inefficient biological light capture with efficient photovoltaics, and using the extracellular electron transport (EET) machinery to deliver this electricity over long distances in microbial cultures and biofilms. This new architecture has the potential to create a better match between the rate of energy delivery and the ability of carbon-fixing metabolism to receive it, allowing the production of enzymatically tailored energy storage molecules and fuels from CO_2_. It is speculated that this hybrid photosynthetic architecture could significantly increase overall energy storage efficiency over natural photosynthesis.

If realized, this ability will allow microbial metabolism to not only be powered by solar electricity, but any renewable electricity source. The nascent field of *microbial electrosynthesis* or simply *electrosynthesis*^36,37^ aims to put biology onto the grid.

The literature review presented here is by no means exhaustive. The reader is referred to other excellent reviews for additional discussion of the literature and topics not covered here^36-39^. This article will focus upon recent work on the molecular basis for the value of the energy conversion efficiency of photosynthesis and consider how these insights relate to the potential conversion efficiency of electrosynthesis. Finally, the article reviews the physical constrains upon electrosynthesis to begin to establish a framework to quantify the overall energy conversion efficiency of this process.

## 2. Introduction to Electrosynthesis

At its core, electrosynthesis aims to substitute the light capture and water splitting machinery of the photosynthetic cell with photovoltaic conversion of light to electricity and electrochemical splitting of water. There are at least three key research and engineering challenges that face the practical implementation of engineered microbial electrosynthesis. The first is to determine the extracellular electron transport (EET) machinery best able to distribute renewable electricity to the optimum number of cells to utilize all of this energy and charge with the highest rate and the least waste. The second challenge is to identify the electron uptake machinery best suited to bring these electrons into the cell and use them to produce intracellular reducing equivalents (NADH, NADPH or ferredoxin) and generate ATP that can then be used to fix CO_2_ and synthesize fuels and energy storage molecules. Lastly, a host or chassis organism, or collection of them, that is best able to assemble the extracellular matrix needed for EET, best equipped to uptake electrons and best suited for CO_2_ fixation and fuel synthesis needs to be identified.

Figure 1 shows a schematic of a photosynthetic cyanobacterium alongside two different electrosynthetic organisms,each utilizing a different electron uptake mechanism that could be used in an engineered electrosynthetic organism.

**Figure 1:**
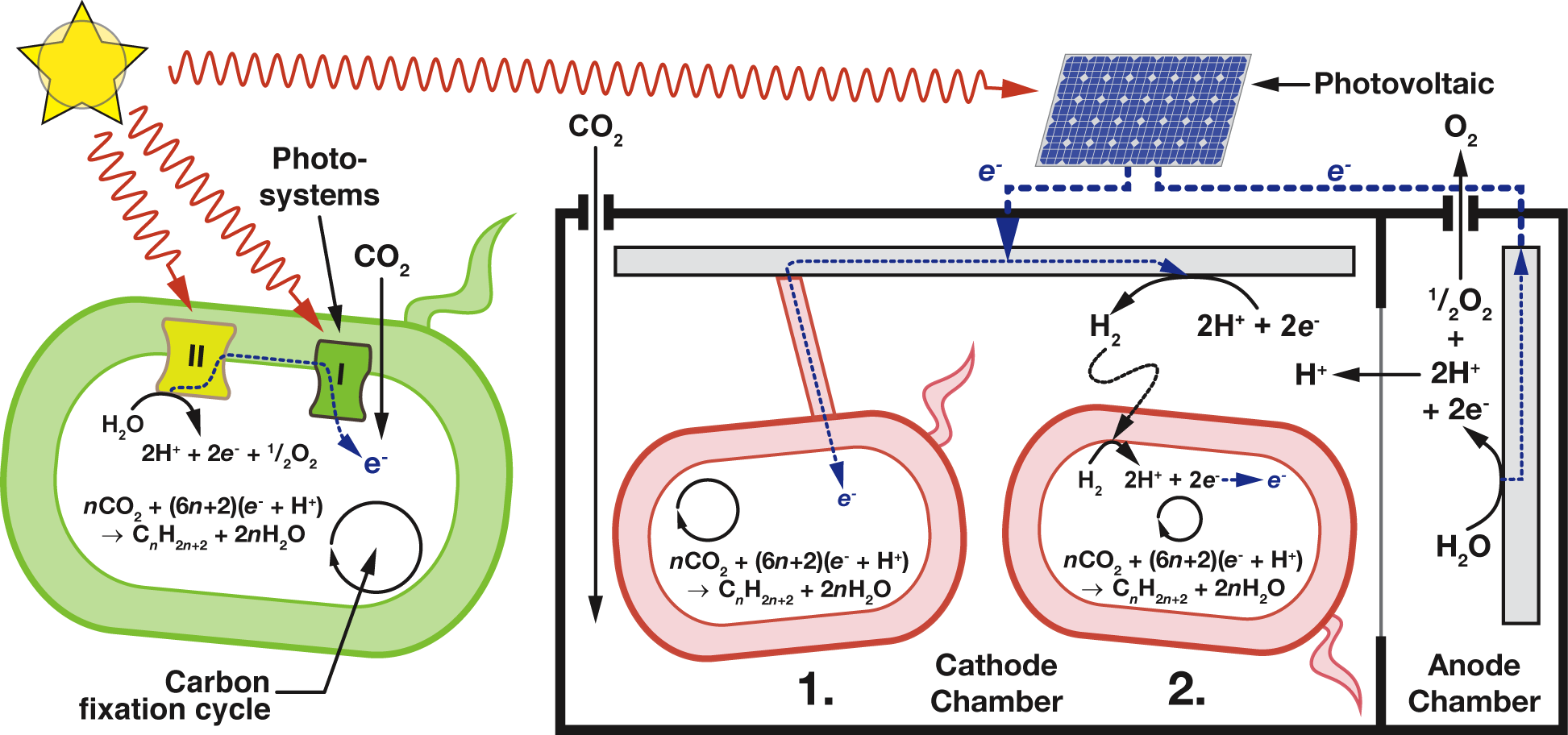
Similarities and differences between photosynthesis and electrosynthesis. On the left is a photosynthetic bacterium, engineered to synthesize alkanes. On the right are two electroactive bacteria engineered to synthesize alkanes inside an electrosynthesis cell. Bacteria 1 (middle) receives electrons through solid matrix conduction (a bacterial nanowire). Bacteria 2 receives electrons through the oxidation of molecular hydrogen that is electrochemically produced at the cathode from protons that are produced by water-splitting at the cell anode.

At the time of writing, there are three possible classes of electrosynthesis host organism. Recent years have seen a wave of renewed interest in naturally occurring microorganisms that specialize in the uptake of electrons from the oxidation of external substrates including metals such as Fe and Mn, to power autotrophic CO_2_-fixing metabolism^40-50^. Recent investigations have demonstrated that these external substrates can be substituted for electrodes, suggesting that an electrochemical cell could drive this metabolism^48,49^. Most notably, several studies have shown that pure cultures of the carbon-fixing organisms *Rhodopseudomonas palustris*^51^, *Sporomusa ovata*^52^ and *Mariprofundus ferrooxydans* PV-1_48_ are capable of uptaking electrons directly from a cathode. The first possible option for the electrosynthesis chassis organism is a version of one of these microbes that has been genetically engineered with biofuel synthesis pathways.

It is important to stress that to date, demonstrations of electrosynthesis have been made almost exclusively in the lab. By contrast, a large number of commercial-scale demonstrations of the cultivation of photosynthetic microorganisms in photobioreactors that permit high cell densities, enriched CO_2_ and mixing to facilitate even light distribution and consequently greatly improved photosynthetic efficiency over that of terrestrial plants have been made^53,54^. That being said, the iron-oxidizing organism and potential electrosynthesis chassis *Acidithiobacillus ferrooxidans* is used as a key component of the copper bio-leaching operation at the Escondida Mine in the Atacama Desert in Chile. With a length of 5 km, width of 2 km and a volume of approximately 1 trillion liters, this bio-leaching heap is arguably the world’s largest bioreactor^44^.

The second option for the chassis organism is to modify an electroactive microbe that specializes in electron outflow rather than uptake. For much of the first decade of the 21^st^ century, most attention was garnered by electroactive organisms that specialize in electron efflux from their metabolism. By depositing electrons derived from catabolism of nutrients onto external substrates such as metals, organisms such as *Shewanella oneidensis* MR-1 and *Geobacter sulfurreducens* are able to respire at high rates under anaerobic conditions^55^. Naturally occurring substrates can be substituted for electrodes, permitting the microbe to power a device called a microbial fuel cell^55^. These extracellular electron transport (EET) processes have been observed to have efficiencies as high as 65%^56,57^. Ross *et al.* demonstrated that the electron outflow organism *S. oneidensis* MR-1 can uptake electrons from an electrode and use them to drive periplasmic reactions^58^. As these organisms are typically heterotrophic, some; most notably *S. oneidensis*; can be easy to culture and genetically engineer. However, this class of microbes do not possess the ability to fix CO_2_. Thus, if one were to be used as the chassis organism, it would require the addition of genes encoding both CO_2_ fixation and fuel synthesis.

The final chassis option is to build an electrosynthetic organism *de novo*. Jensen *et al.* transferred the Mtr electron outflow system from *S. oneidensis* to *E. coli* and demonstrated its *in vivo* functionality^59^. Complementary success in transporting genes encoding the 3-hydroxypropionate (3HOP) carbon fixation cycle to *E. coli*^60^, along with numerous demonstrations of fuel and fuel precursor synthesis in *E. coli* raise the possibility of constructing an electrosynthetic strain of *E. coli*.

In addition to the choice of host organism, two broad classes of EET mechanisms are available for the connection of biological metabolism to an electrode. In recent years, considerable attention has begun to focus upon mechanisms that permit the transport of renewable electricity to the cell rather than from it^61^.

The EET mechanism that is perhaps most synonymous with microbial fuel cells, and initially with electrosynthesis, are conductive pili (sometimes called bacterial nanowires)^62^ that transport electrons between the cell surface and external substrates at high rates. This mechanism is sometimes referred to as *solid matrix conduction*. A microbe utilizing this mechanism is shown in case 1 in figure 1. Specialized inner and outer membrane spanning complexes can connect with this solid matrix and provide a conductive path for electrons from metabolic reactions in the cytoplasm and periplasm of the cell to sites a long distance from the cell surface^55,63^. It is possible that solid matrix conduction can transfer electrons over centimeter length scales^64^. This mechanism of EET is best known from microbial fuel cells^65^ and is speculated to have the most promise given the high rates of electron transfer observed in microbial fuel cells^37^.

A second option for electron transport (case 2 in figure 1) is a soluble electron carrier (or mediator) that is electrochemically reduced at the cathode, transported by diffusion to the microbe and oxidized. Of all the strategies reviewed here, this has enjoyed the most success in engineered electrosynthesis. Considerable recent attention has focused on the use of H_2_ as a mediator. Torella *et al.* used H_2_ produced by a low cost anode to power CO_2_ fixation and propanol production by the H_2_-oxidizing bacterium *Ralstonia eutropha*^66^. Li *et al.* used electrochemical reduction of CO_2_ to formate to drive butanol production in *R. eutropha*^67^.

An extensive body of work has demonstrated the use of low potential synthetic soluble mediators to transfer electrons from an electrode to *in vivo* redox processes and increase the yield of more reduced fermentation products. Rao *et al.* showed increased alcohol production in *Clostridium acetobutylicum* in response to reduced methyl viologen exposure^68^. Kim *et al.* demonstrated the use of methyl viologen to increase butanol production in *Clostridium acetobutylicum*^69^. Park *et al.* showed that exposure to neutral red will increase fumarate reduction by *Actinobacillus succinogenes*^70^. Lovely *et al.* demonstrated that electrons supplied by reduced anthraquinone-2,6-disulfonate (AQDS) could allow growth of *Shewanella alga* on a carbon source normally too oxidized for use^71^. Most recently, Steinbusch *et al.* used methyl viologen to increase acetate reduction in mixed cultures^72^. Rosenbaum *et al.* made an extensive review of biologically synthesized mediators ^61^. Additionally, there have been numerous demonstrations of the use of synthetic mediators to enhance current production in microbial fuel cells^65^.

## 3. The Importance of Energy Storage Efficiency

The benefits of electrosynthesis, replacing inefficient photosystems with efficient photovoltaics, are qualitatively apparent. However, a quantitative understanding of the efficiency losses in photosynthesis are necessary to fully assess if hybrid approaches can successfully address them. In addition, if electrosynthesis can indeed address these inefficiencies, an understanding of the energy losses in the process are needed in order to reduce them and allow the approach to make full use of its potential.

This article makes no pretense of fully reviewing the extensive and excellent body of literature on the inefficiencies of photosynthesis. Three goals of this article are to begin to quantitatively compare electrosynthesis and photosynthesis, to begin to assess if electrosynthesis can address the shortcomings of photosynthesis, and to inspire a framework for the quantitative assessment of the inefficiencies of electrosynthesis.

While photovoltaics with electrochemical storage, photosynthesis and hybrid photosynthesis all differ substantially, they can all be compared on a common scale: overall storage efficiency, *ηo*, that relates how much retrievable chemical energy, 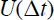, can be stored per unit area in a given time interval, Δ*t*, relative to the solar radiation arriving at ground level in that time, 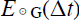,

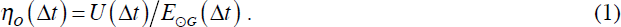

To illustrate potentially acceptable energy storage efficiencies, imagine if all of the US transportation energy requirements were supplied by solar energy capture. In 2013, the last full year for which data is available, the United States transportation sector consumed 28.8 EJ^21^, corresponding to an average instantaneous rate, 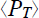, of 0.91 TW. In order to capture this, a minimum land area,

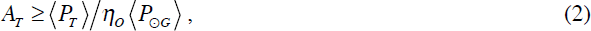

is required, making the assumption that no energy losses in conversion of the storage chemical to transportation fuel. The results of this formula are compiled in table 1 for the annually averaged solar insolation at ground level across day and night 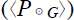 at a latitude of 30° N of2248210 W_29_ for a range of efficiencies representing a variety of energy capture and storage scenarios.

**Table 1.** Minimum land areas required to capture 0.91 TW of solar power for a range of capture and storage efficiency scenarios. Columns 3 to 7 report the land area as a fraction of the special use-; crop-; forest-; grass-, pasture- and range-; land and total land area of the lower 48 United States^118^.

| Overall Efficiency ($\eta_o$) (%) | Minimum Land Area ($m^2$) | Special Use including Parks and Wildlife | Cropland | Forests | Grass, Pasture and Range | Total Area of Lower 48 States | Comment |
| --- | --- | --- | --- | --- | --- | --- | --- |
| 0.1 | $4.3 \times 10^{12}$ | 6.30 | 2.6 | 1.9 | 1.7 | 0.47 | Pessimistic estimate for the overall storage efficiency of photosynthesis. |
| 0.25 | $1.7 \times 10^{12}$ | 2.50 | 1.0 | 0.74 | 0.70 | 0.19 | Brenner's estimate for the overall efficiency of photosynthesis<sup>29</sup>. |
| 0.5 | $8.7 \times 10^{11}$ | 1.30 | 0.52 | 0.37 | 0.35 | $9.3 \times 10^{-2}$ | |
| 0.75 | $5.8 \times 10^{11}$ | 0.84 | 0.35 | 0.25 | 0.23 | $6.2 \times 10^{-2}$ | |
| 1 | $4.3 \times 10^{11}$ | 0.63 | 0.26 | 0.19 | 0.17 | $4.7 \times 10^{-2}$ | Realistic upper estimate on photosynthetic efficiency in the real world<sup>30</sup>. |
| 1.5 | $2.9 \times 10^{11}$ | 0.42 | 0.17 | 0.12 | 0.12 | $3.1 \times 10^{-2}$ | |
| 3 | $1.4 \times 10^{11}$ | 0.21 | $8.7 \times 10^{-2}$ | $6.2 \times 10^{-2}$ | $5.8 \times 10^{-2}$ | $1.5 \times 10^{-2}$ | Algal culture in photobioreactor<sup>54</sup>. |
| 14 | $3.1 \times 10^{10}$ | $4.5 \times 10^{-2}$ | $1.8 \times 10^{-2}$ | $1.3 \times 10^{-2}$ | $1.3 \times 10^{-2}$ | $3.3 \times 10^{-3}$ | Commercial PV coupled with H<sub>2</sub> storage<sup>30</sup>. |
| 20.8 | $2.1 \times 10^{10}$ | $3.0 \times 10^{-2}$ | $1.3 \times 10^{-2}$ | $8.9 \times 10^{-3}$ | $8.4 \times 10^{-3}$ | $2.2 \times 10^{-3}$ | Highest efficiency Si PV coupled with H<sub>2</sub> storage<sup>24</sup>. |
| 36.8 | $1.2 \times 10^{10}$ | $1.7 \times 10^{-2}$ | $7.1 \times 10^{-3}$ | $5.1 \times 10^{-3}$ | $4.8 \times 10^{-3}$ | $1.3 \times 10^{-3}$ | Highest efficiency multi-junction PV coupled with H<sub>2</sub> storage<sup>24</sup>. |

It is important to stress the inverse relationship between minimum area and efficiency. This relationship has the consequence that apparently insignificant changes in efficiency, especially at the low end of the range, can result in large changes to the minimum land requirement.

When the land areas calculated in table 1 are converted to fractions of available land area in the US, they range from worrisome to barely worthy of note. It should be stressed that no matter what technology is used to capture solar power, low efficiencies are doubly problematic as they not only occupy more land, but they increase the financial and energy costs of collecting stored energy and transporting it for processing and distribution.

## 4. Estimating the Energy Conversion Efficiency of Photosynthesis

### 4.1 Definition of Photosynthetic Efficiency

Photosynthesis is the most relevant point of comparison for electrosynthesis given the success of energy crops and as it shares a considerable amount of molecular machinery with electrosynthesis. This comparison allows us to assess where the inefficiency of photosynthesis arises, and if electrosynthesis could successfully address these drawbacks. Additionally, this analysis allows the opportunity to assess the performance of some of their shared molecular machinery in the real world. Finally, similarities between the two processes could allow innovations in one area to be rapidly transferred to the other.

Monteith introduced the concept of genetic yield (Y)^73^ (later adapted by Zhu *et al*.^74^) that quantifies the extractable quantity of energy stored under ideal conditions of growth per unit area of land over a span of time Δ*t* (typically a growing season). Genetic yield is typically expressed as the product of a series of sub-efficiencies,

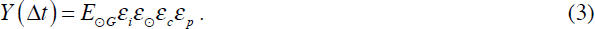

The interception efficiency, 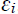, is the fraction of light intercepted by leaves in the canopy. 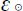 is the fraction of the solar spectrum, often called the photosynthetically active radiation (PAR) portion that can be absorbed by the leaf. 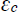 is the photosynthetic conversion efficiency and is the fraction of the intercepted photosynthetically active radiation that is stored as biomass. The product 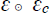 is sometimes referred to as the full spectrum photosynthetic conversion efficiency. Finally, 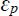 is the partitioning efficiency and relates the portion of energy in biomass that is harvestable.

During composition of this article, it has been intellectually useful to express *Y* as the integral of the product of the sub-efficiencies. This re-casting emphasizes the importance of the time and solar power dependence of these factors.

The position of a plot of land changes with respect to the Sun, with both time of day and season. This relative position change affects the intensity, 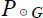 spectral content and angle of the incident light. Changes in spectral content affect spectrum utilization efficiency, 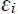; while changes in angle affect interception efficiency, 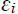. Most importantly, the changes in the delivered power have the potential to affect the conversion efficiency, 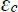. Thus, it is useful to re-cast the genetic yield as an integral that reflects its dependence upon time; solar intensity; latitude, *φ* and longitude, *λ* of the plot of land on the Earth’s surface,

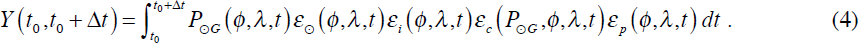

The efficiency of photosynthesis can be calculated using equation 1 and by equating *Y* to U,

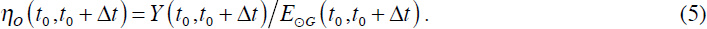

The concepts of efficiency in relation to photosynthesis have at times been used inconsistently in the literature, leading to a range of reported values, sometimes referring to different physical quantities^75^. It is useful to clarify these concepts and to make a simple global scale estimate of the real-world efficiency of photosynthesis. Given the ready availability of data for the land, this analysis is restricted to terrestrial photosynthesis.

The total energy stored per square meter of land, *U_b_*, can be estimated using satellite measurements of net primary productivity (NPP)^76^ which quantifies the net mass, *mc*, of carbon absorbed by a unit area of land at a position (φ, λ) on the Earth’s surface. Stored energy can be estimated by assuming that all fixed carbon is converted to as wood-like substance. Dry wood has an energy content, *u_w_*, (lower heating value) of ≈ 19 kJ/g^77^ and a carbon content, *cc*, ranging from ≈ 47 to 55%^78^. Thus, the estimated energy stored in biomass per square meter,

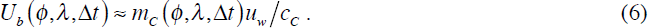

Satellite data also gives the average incident solar flux at ground level 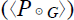 and can be used to calculate the total amount of solar energy, 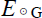, arriving at ground level per square meter of land during Δ*t*.

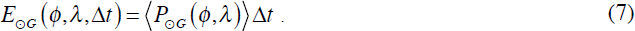

With this in hand, one can readily estimate the overall photosynthetic efficiency. It is important to stress that not all of this energy might be recoverable, but by assuming that 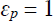, it is possible to place an upper bound on the overall efficiency defined in equation 1,

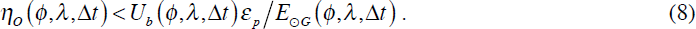

Estimates for overall photosynthetic efficiency during June of 2014, the last peak of NPP in the northern hemisphere on record, are plotted on a map the world in figure 2A. This snapshot of carbon uptake was chosen in order to observe the highest possible storage rates, and hence efficiencies, and permit a more direct comparison with theoretical estimates of maximum photosynthetic efficiency. Overall efficiencies range from negative, indicating more respiration than carbon fixation, to 1% in areas of peak forest growth in northern Europe, North America, Russia, northern Asia and the Amazon. This estimate, made at the time of annual peak plant growth on Earth, agrees roughly with the estimate of the upper bound on real-world photosynthetic conversion efficiency noted by Blankenship *et al*.^30^.

**Figure 2:**
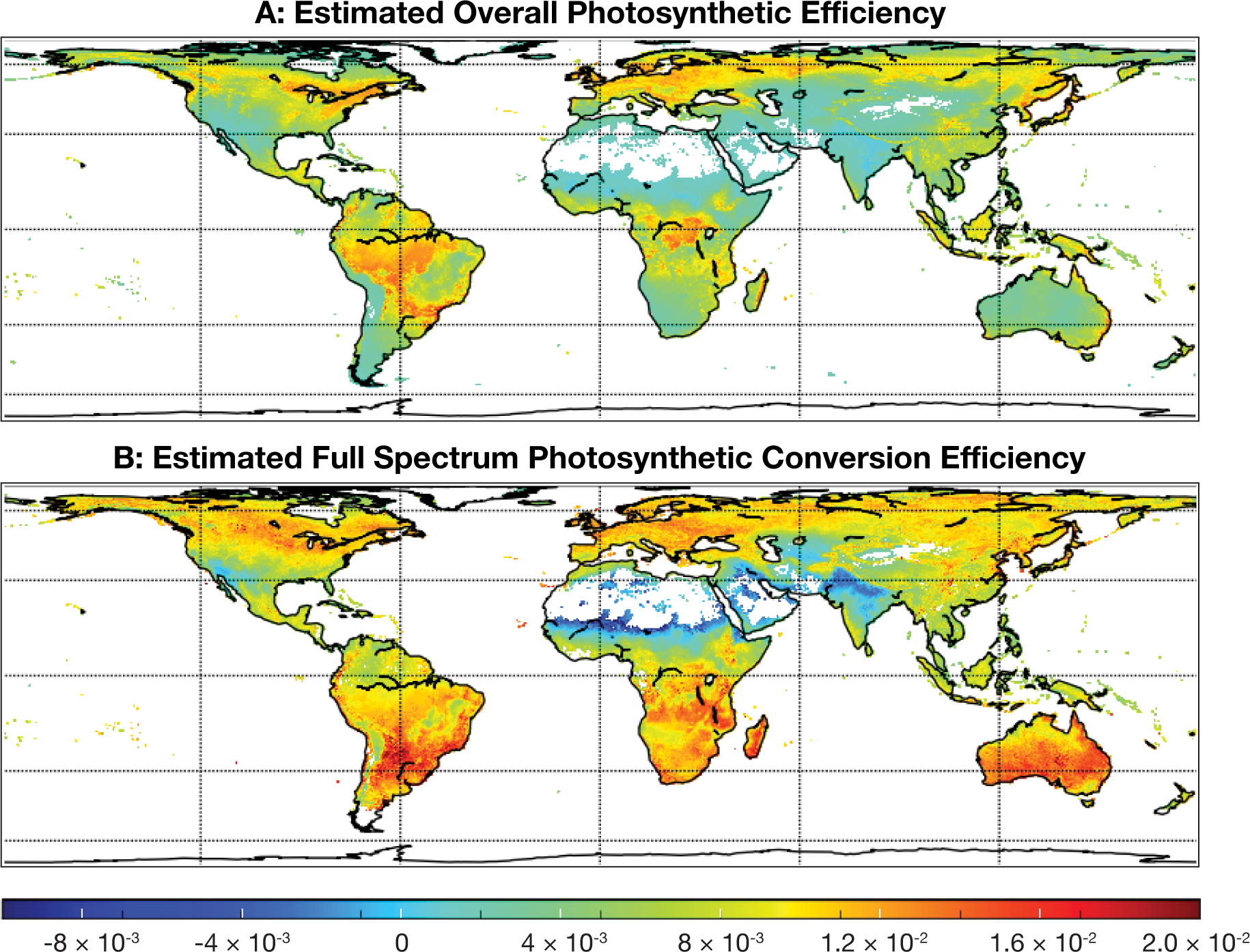
A: Estimateci overall photosynthetic efficiency. Β: The overall estimated efficiency normalized by estimated interception efficiency. The false color bar denotes the efficiency as a fraction of 1.0 and the color bar is shared between plots. Net primary productivity data were derived from measurements by the MODIS instrument on the Terra satellite^76^. Insolation, NPP and Leaf Area Index data are available at^120-122^.

It is important to stress that while overall efficiency is an important quantity, normalization provides additional insight into photosynthetic efficiency. An analogy is useful to understand this: consider two solar photovoltaic farms, one with closely spaced modules and another with distantly spaced modules. The first will collect more solar energy per unit time, but this fact will simply reflect the spacing of the modules chosen by the designer of the farm, not the intrinsic conversion efficiency of the modules or the cells that make up the modules that are recorded in a lab test and reported in the literature. In addition, panel orientation along with damage, dust and other real world insults will also further increase the difference between real-world and lab efficiency.

Given that the leaves of plants in the real world often suffer from damage due to hail, frost, fire or lodging (stem breakage) and are sometimes planted at sub-optimal spacings, it is useful to normalize the overall photosynthetic efficiency by the interception efficiency. This can be estimated using satellite measurements of leaf area index (*L*) which quantifies the amount of leaf cover per square meter of land, and the canopy extinction coefficient, *k*^79^,

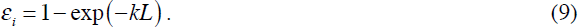

For planophile canopies with horizontal leaves, *k* ≈ 0.9, while for erectophile canopies with vertical leaves, *k* ≈ 0.3_79_. For a canopy with a random leaf orientation and inclination, *k* ≈ 0.5^80^. Thus,

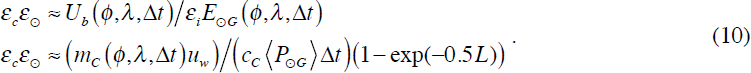

These estimates are plotted in figure 2B. Normalized full spectrum photosynthetic conversion efficiencies peak at over 1.6% in some parts of North America, and northern Europe. Interestingly, normalized conversion efficiency is also high in the Outback of Australia, even though the overall efficiency is extremely low because of minimal leaf cover.

While the overall energy conversion and full spectrum conversion efficiencies of photosynthesis across much of the globe are low, resulting in rather large land area requirements for any photosynthetic energy storage scheme, meaningfully higher rates are possible both in principle and have been achieved in practice.

Overall full season efficiencies well in excess of 1% are possible in the the most productive growth regions in the world such as the US Midwest and the United Kingdom^33^. These efficiencies are in large part attributable to the use of crop strains bred for good canopy formation along with highly developed planting techniques and irrigation, fertilizer and pesticide use which contribute further to canopy formation and high interception efficiencies.

At maximum growth rates, under the temperate, but relatively low insolation levels of England, C_3_ plants in the field can operate at maximum overall conversion efficiencies as high as 3.5%, while C_4_ plants attain a conversion efficiency of ≈ 4.3%^81^. When averaged over a full growing season, these efficiencies drop to ≈ 2.4% for C_3_ plants including sugar beets, barley potatoes and apples^73^ and ≈ 3.4% for C_4_ plants such as *Miscanthus χ giganteus*^82^. Algae grown in bubbled photo-bioreactors can achieve annually averaged efficiencies of ≈3%^54^.

Zhu *et al.*^74,83^ conducted a survey of the loss mechanisms of photosynthesis, and calculated upper bounds for the full spectrum conversion efficiency 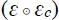 of C_3_ and C_4_ photosynthesis. The sources of energy losses identified by this breakdown highlight the potential gains in efficiency from hybrid photosynthesis approaches such as electrosynthesis and potential avenues for improvement of photosynthesis. A modified summary of the energy losses in C_4_ photosynthesis is plotted in figure 3.

**Figure 3:**
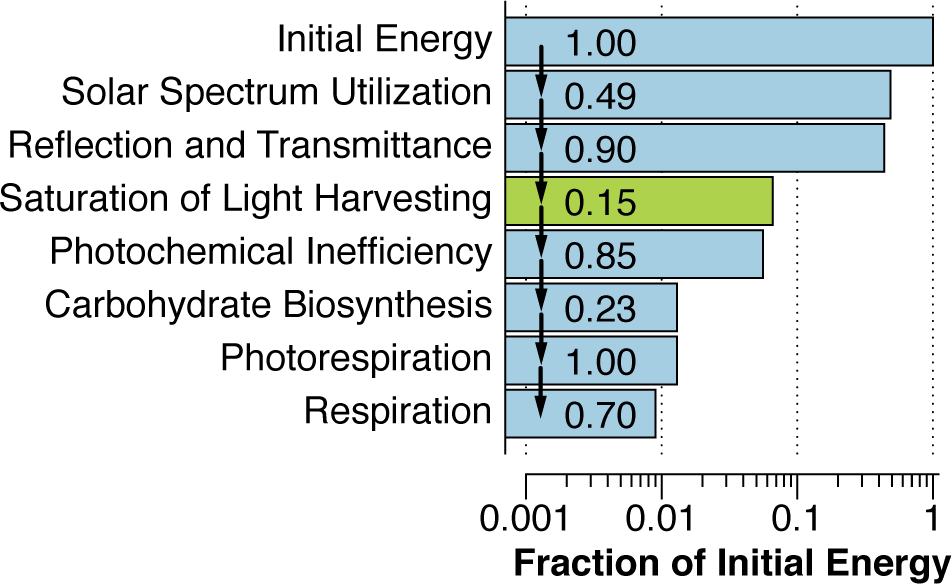
Model of energy losses in C_4_ photosynthesis built upon a model by Zhu *et al?*^4^ with the addition of an average 85% energy loss due to saturation of light harvesting (green bar). The cumulative efficiency is indicated on the horizontal axis, while the efficiency of each step is shown in each bar.

### 4.2 Solar Spectrum Utilization - 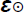

The majority of terrestrial plants utilize a portion of the solar spectrum that spans from ≈ 400 to ≈ 740 nm and contains only ≈ 49% of the power present in the AM1.5 global tilt spectrum^84^. Put another way, for each 1000 J of solar energy incident upon a plant, only 487 J are available for photosynthesis^83^. This utilization window is set by chlorophyll a and b, the primary pigments that compose the light harvesting complexes of photosystems I and II. By contrast, Si solar photovoltaics are able to access wavelengths out to ≈ 1130 nm, permitting usage of up to ≈ 77% of the power content of the solar spectrum. In addition, it is estimated that leaves reflect ≈ 10% of the incident photosynthetically active radiation, denying plants another 49J^83^.

### 4.3 Conversion Efficiency - 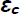

Following photon absorption by the light harvesting complexes, de-excitation is required prior to absorption by the reaction center chlorophylls of Photosystems I and II whose absorption spectra peak at 700 and 680 nm respectively. This results in the loss of another ≈ 66 J of the initial 1000 J, leaving only 372 J. C_3_ carbon fixation deposits ≈ 34% of the solar energy remaining after de-excitation into glucose, leaving ≈ 126 J of the initial energy. A further 61 J is lost to photorespiration under present day CO_2_concentrations (380 ppm) and at a temperature of 30° C, while an additional ≈ 19 J is lost to respiration. This leaves 46 J of the original 1000 J, for a final conversion efficiency of 4.6%.

In C_4_ photosynthesis, 287 J are lost in fixing CO_2_ to glucose. However, the additional energy expenditure of carbon concentrating mechanism comes with the benefit of eliminating photorespiration losses. An additional 25 J are lost to plant respiration, leaving 60 J of the original 1000 J, for a final conversion efficiency of 6.0%. Given that cyanobacteria utilize carboxysomes for carbon concentration, the upper bound on photosynthetic efficiency for this class of organisms is likely to resemble that for C_4_ photosynthesis. A summary of these losses for C_4_ photosynthesis is plotted in figure 3.

There is a growing consensus that plants in the field do not achieve these upper bound efficiencies because photosynthesis has not been optimized by evolution to gather and store the largest amount of light energy. While the rate of photosynthesis rises linearly with illumination at low light levels, this rate plateaus at between 10% of the maximum solar flux for shade plants and 20% for sun plants^29^.

A simple calculation indicates that the supply of photons to a photosynthetic cell can quickly outstrip photon demand for carbon fixation. The photon interception cross-section of the cell can be calculated from the radius of gyration of the cell. Assuming a cylindrical cell like *E. coli* with a diameter *d* of ≈ 1.3 μm and a length *l* of ≈ 3.9 μm^85^, the radius of gyration,

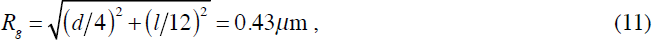

while the effective interception cross-section,

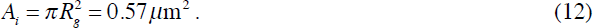

Assuming as before that only 48.7% of the energy content of the solar spectrum is useable and that the average energy per mole of PAR photons is 205 kJ^83^, the delivery rate of PAR photons to the cell,

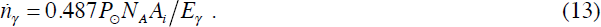

This rate is plotted for a range of solar intensities, ranging from 0 to 1030 Wm^-2^ (the peak solar power at ground level) in figure 3.

The photon demand of the cell can be calculated by estimating the amount of RuBisCO, the primary carboxylating enzyme in the Calvin cycle, hosted by the cell and the catalytic rate of this enzyme. Assuming a dry weight for the cell of 433 fg (Bionumbers ID number 103892^86^) and that protein constitutes 52.4% of this dry weight (ID number 101955^86^), the total dry weight of protein in the cell,

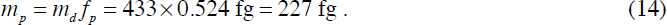

Leaves maximize carbon fixation by maximizing synthesis of RuBisCO: this enzyme constitutes over 50% of leaf soluble protein^83^, making it probably the world’s most abundant enzyme^87^. With a molecular weight of 65 kDa per catalytic unit (ID number 105007), the estimated number of RuBisCO catalytic units in the cell,

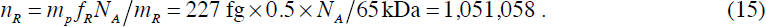

Assuming a photon demand of between 8 (for C_3_ photosynthesis) to 12 (C_4_)photons per carbon fixation, the total photon demand rate,

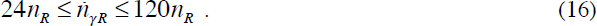

The upper bound of this estimate is plotted in figure 4 and suggests that the Calvin cycle can only usefully utilize between ≈; 5 to 15% of the maximum solar flux, close to the point at which photosynthesis typically saturates^29^.

**Figure 4:**
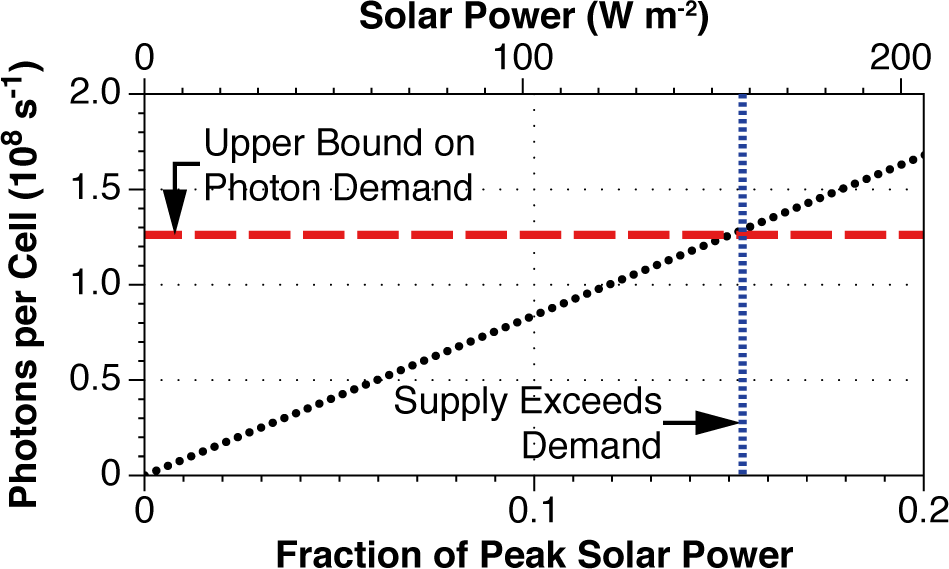
Plot of photosynthetically active photon supply incident upon a cell with a light interception cross section of ~ 0.6 μιη^2^ as a function of the fraction of peak solar power at ground level (~ 1030 Wm^-2^). The dashed horizontal red line shows an upper estimate of photon demand for carbon fixation under ambient CO_2_ concentrations. Photon supply begins to exceed demand at ~ 0.15 of the peak solar power (~ 158 Wm^-2^).

The low catalytic rate of RuBisCO and light harvesting saturation suggests a fundamental mismatch between the size of the photosynthetic cell and its area. Rather than allowing this light to pass through, plants typically dissipate this excess light as heat, denying it to competitors by shading. Gains in photosynthetic rate by increasing the amount of RuBisCO per leaf seem largely to have been exhausted as this protein already constitutes ≈ 50% of leaf soluble protein and it is doubtful that more could be added without health consequences to the leaf or the use of undesirable quantities of fertilizer^83^.

To illustrate the effect of this saturation effect on photosynthetic efficiency, an extra loss step has been added to the analysis by Zhu *et al*.^83^ in figure 3. It should be stressed that the saturation effect is dependent upon light intensity, so its effect will vary with time of day and season. However, an average saturation loss of 85% does account for much of the difference between maximum photosynthetic rates and observed rates in the field. This analysis indicates that a significant portion of the inefficiency of photosynthesis occurs due to limited access to the full solar spectrum, saturation of light harvesting saturation and inefficiency of photon capture. Were photosynthetic cells able to fix more carbon, it is possible that the overall conversion efficiency of photosynthesis could rise substantially.

## 5. Research Challenges in Electrosynthesis

The significant energy losses that occur during the light harvesting and capture steps in photosynthesis support some of the attraction of electrosynthesis. Rather than increasing the carbon fixation capacity of an individual cell, electrosynthesis aims to increase the number of cells that are able to access the reducing equivalents produced by sunlight. In order to validate this attraction, much more needs to be learned about the molecular mechanisms underlying electrosynthesis.

### 5.1 Bringing Electrons to the Cell Surface

The two broad classes of extracellular electron transport were shown in the introduction and figure 1. Soluble mediator driven extracellular electron transport is exemplified by hydrogen-mediated electron transport and is shown in case 1 in figure 1, while solid matrix conduction is shown in case 2 in figure 1. Both hydrogen-mediated transport and solid matrix conduction offer their own sets of attractions and drawbacks.

As discussed in the previous section, much of the overall inefficiency of photosynthesis derives from the large mismatch between the carbon-fixing capacity of a photosynthetic cell, its light capture ability and the available light. This architectural mismatch forces photosynthesis, under peak illumination, to forfeit close to 90% of the opportunities it has to generate reducing equivalents.

EET offers the opportunity to distribute solar-generated reducing equivalents over a volume of carbon-fixing cells sufficient to make full use of them. The architecture used to host these cells will be enormously influenced by the molecular mechanism used for distribution.

To understand how the two classes of EET can each differently influence the architecture of an electrosynthesis reactor it is useful to estimate the electron consumption needs of a single carbon-fixing and fuel synthesizing cell and the total reducing equivalent generation capacity of a unit area of land.

The synthesis of butanol (an infrastructure compatible biofuel) requires a minimum of 4 CO_2_ molecules (*n_cb_*) and 24 electrons (*n_eb_*)^37^. Assuming a cell of the same dimensions and containing the same number of RuBisCO molecules as in equation 15 and a turnover rate range from 3 to 10 CO_2_ s^-1^, the electron rate requirement,

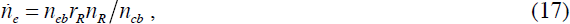

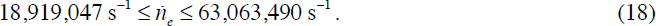

The total reducing equivalent generation potential of a unit area of land can be estimated by counting the total number of photons incident upon that land with an energy exceeding that needed to split water and reduce protons to H_2_. This photon count is estimated by numerically integrating the AM1.5 global tilt spectrum^84^ from 400 nm to 1008 nm (corresponding to an energy of 1.23 eV) and counting photons in each wavelength bin by dividing the power by the energy of each photon at that wavelength. The photon delivery per square meter of ground,

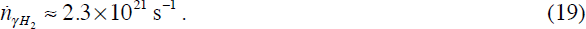

This delivery rate corresponds to an effective potentially available current density,

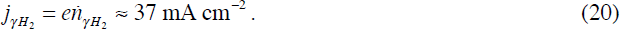

#### 5.1.1 Hydrogen-mediated Electron Transport

Figure 5A shows a model for electron uptake from H_2_ oxidation utilized by *R. eutropha*^88^. In this model, the organism uses its membrane bound NiFe-hydrogenase (MBH) to supply electrons to its electron transfer chain for ATP generation. The soluble hydrogenase (SBH) is used for the direct reduction of NAD+, which is then used to generate NADPH via a transhydrogenase reaction^89^. This NADPH is used for subsequent biosynthetic reactions, including carbon fixation.

**Figure 5:**
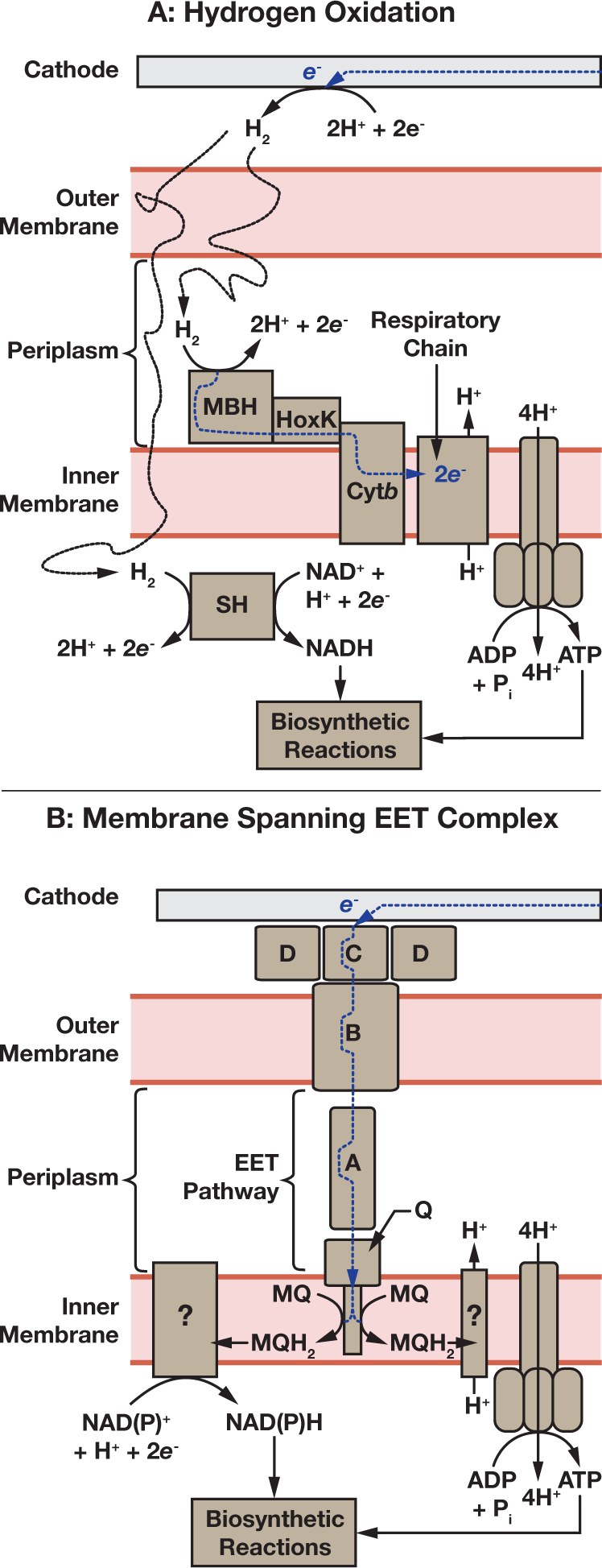
Model of electron uptake processes. A: model of electron uptake by hydrogen oxidation by *Ralstonia eutropha* adapted from^88^. MBH is the Membrane Bound Hydrogenase, and SH is the Soluble Hydrogenase. B: A general model of electron uptake by a neutrophilic organism employing a periplasm-spanning extracellular electron transport (EET) pathway such as Mtr, Pio or Mto. For *Shewanella oneidensis*,A denotes MtrA, B - MtrB, C - MtrC, D - OmcA and Q - CymA.

Four outstanding features of H_2_-mediated EET offer the potential for high solar to liquid fuel conversion efficiencies. This approach is aided at the outset by the high efficiencies of commercial electrolyzers that are capable of storing electrical energy as H_2_ with efficiencies approaching 80%^30^. Blankenship *et al.* estimate that while the energy losses due to potential mismatches in present day photovoltaics and electrolyzers can be as high 30%, these losses can be greatly reduced^30^. Secondly, due to its extremely low viscosity, H^2^ can be transported over very long distances with minimal energy loss^90,91^. This feature is particularly attractive as it allows the possibility for H_2_ generation and carbon fixation to be spatially separated and independently optimized.

Thirdly, potential mismatches within H_2_-oxidizing organisms are already limited. The redox potential of H_2_/2H^+^ + 2*e*_-_ is -0.42 V relative to the Standard Hydrogen Electrode (SHE), whereas for NAD+ + H^+^+ 2*e*^-^/NADH it is -0.32 V, representing only ≈ 8% of the energy of a 1.2 eV photon.

Finally, the protein requirements of H_2_ oxidation are low relative to carbon fixation. Given that each H_2_ molecule delivers two electrons to the cell, and a hydrogenase turnover rate r/, the number of hydrogenase enzymes needed to supply the electron requirements of fuel synthesis,

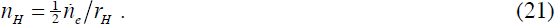

For example, the *Chromatium vinosum* NiFe uptake hydrogenase^92^ has a turnover rate between 1500 and 9000 H_2_s^-1^. Thus,

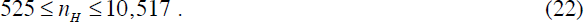

Given a molecular weight, *m*_H_ of 94 kDa for the *C. vinosum* hydrogenase, the total mass of hydrogenase needed to satisfy this electron uptake,

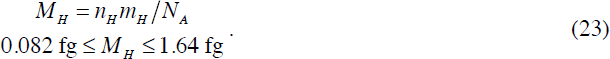

This mass constitutes between only 0.04% and 0.7% of the total dry protein mass of the cell (equation 14). The low protein demands of this uptake method certainly recommend this approach and could be readily increased should improved carbon-fixation ability become available.

A commonly expressed concern regarding H_2_-mediated electron transport is the electron delivery rate limitation imposed by the low solubility of H_2_ in water (relative to CO_2_ for instance). It is not apparently obvious that this limitation need have a detrimental effect on overall system efficiency in all cases, but its effect upon overall system architecture could be profound and is worth examining. This can be demonstrated by considering the constraints that diffusional transport poses on system geometry. Consider a volume of water, *V*, containing cells at a density, *ρ_c_*, of 10_12_ per liter. The total H_2_ demand of these cells,

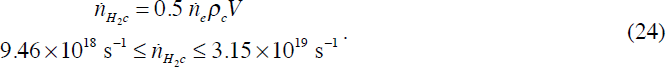

Fick’s first law can be used to calculate the size of the concentration gradient necessary to produce this flux ^93^. The rate of diffusional particle flux into the volume *V* through a planar face of area *A* is equal to the concentration gradient normal to that plane multiplied by the diffusion coefficient of the particle species,

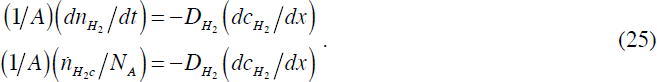

The H_2_ concentration at this plane is equal to the maximum solubility of H_2_ under the conditions of the system. Over the height of the cuboid, Δ*x*, the H_2_ concentration drops to 0 due to consumption by the cells. Thus,

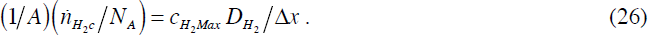

For a system in which H_2_ is supplied through the headspace above the liquid volume, Henry’s Law can be used to calculate the H_2_ concentration at the supply face of the cuboid,

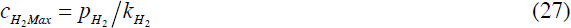

as a function of the partial pressure of H_2_ in the headspace, *p*_H_2__ and the solubility constant for H_2_ in water, *k*_H_2__. Thus, the necessary aspect ratio of the liquid,

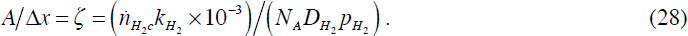

While the surface area and height,

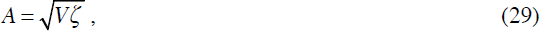

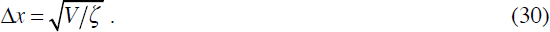

This geometry results in an effective current density,

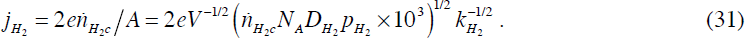

*R. eutropha* is typically grown under an atmosphere containing H_2_, O_2_ and CO_2_ at a ratio of 8:1:1 ^89^. At the laboratory-scale, the H_2_ pressure in the headspace is typically restricted to 5% or less of a total of 1 atmosphere in order to reduce the risks of H_2_ explosion^89^. Under these conditions a 1 L volume is restricted to a geometry of,

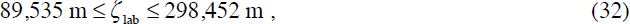

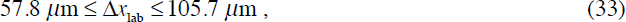

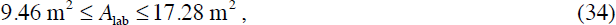

leading to an effective current density of,

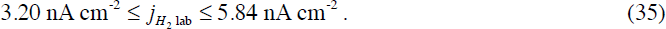

At industrial-scales, it is possible that these restrictions on H_2_ concentration could be lifted. For a headspace with a total pressure of 1 atmosphere containing 80% H_2_, 10% O_2_ and 10% CO_2_,

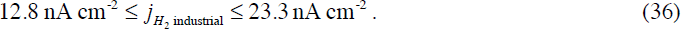

Both of these current densities are 6 orders of magnitude lower than the potentially available solar current density calculated in equation 20. In order to make full use of the light incident upon a unit area of ground, one would need to construct a reactor with an effective area of almost 1 million times greater. This is in itself not intrinsically a problem, but it may require creative engineering to maintain energy conversion efficiency, minimizing H_2_ loss, maintaining an acceptable level of safety and mitigating the effects of proton consumption due to fuel synthesis on solution pH. One concept is the hollow fiber gas reactor designed by Worden and Liu^94^, in which *R. eutropha* biofilms are immobilized on the surface of hollow fibers. A high H_2_ atmosphere is maintained on the exterior of the fiber while a high O_2_ atmosphere is maintained on the interior. This permits packaging of high surface areas within a reasonable volume while affording the culture access to both gases but minimizes mixing and thus circumvents some of the safety concerns of containing both within the same vessel^89^.

The recycled-gas culture system designed by Schlegel and Lafferty^95-97^ increases H_2_ transfer rate by stirring and uses elaborate H_2_ storage system to minimize gas losses.

#### 5.1.2 Solid Matrix Conduction

An alternative to H_2_-mediated electron transport is solid matrix conduction (case 2 in figure 1) in which electrical current is transferred through conductive pili inside a biofilm that is attached to an electrode surface. A notable example of the high electron transfer rates achievable by this method is a current density of 456 μA cm^-2^ recorded flowing from a *Geobacter sulfurreducens* film by Nevin *et al*.^98^. Other observations from *Geobacter* films fall into the range of 8 μA cm^-2^^99^ to 222 μA cm^-2^ ^100^. These current densities are between 3 and 4 orders of magnitudes greater than those predicted for H_2_-mediated transport in equations 35 and 36 and only 2 to 3 orders of magnitude lower than the maximum potential current density that could be extracted from sunlight in equation 20.

Solid matrix conduction has also been demonstrated for electron uptake. However, in contrast, current densities observed for the uptake of electrons have thus far been lower than those seen in outflow and range from 1.5 μA cm^-2^ ^51^ to ≈ 10 μA cm^-2^ ^48^. These current densities are between 4 χ 10^3^ and 3 χ 10^4^ lower than the maximum potential solar generated current density (equation 20).

It is useful to estimate the maximum electron uptake current density that might be achieved by solid matrix conduction given sufficient biological and reactor engineering. The current density needs of a single cell can be made by assuming that electrons are absorbed over the entire surface area of the cylindrical bacteria, *A_cs_*,

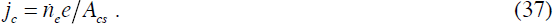

Given that microbes in the film will be separated by extracellular matrix and that cells in each layer will only occupy a fraction *f_c_* of the layer, the current density demands of a single layer of cells,

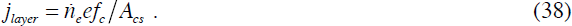

Similarly, each layer of cells, with a height *d*, will be separated from the ones above and below by a distance *h_gap_*. For a film of total height *h_fim_*, the number of layers will be,

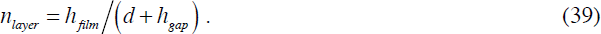

Thus, the total current density of the film,

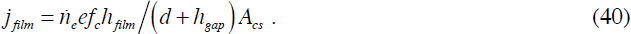

Assuming that that no more than 25% of the area of a layer^101^ is filled with cells of diameter *d* of ≈ 1.3 μm and length *l* of ≈ 3.9 μm, the maximum current density demand that could be expected per layer

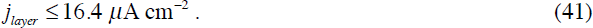

Assuming that each layer of cells is separated by the cell diameter *d*, a film height of ≈ 50 μm^98^ will draw a current density,

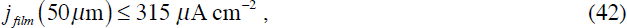

and for a film of 1 cm thickness^64^ the estimated current density requirement,

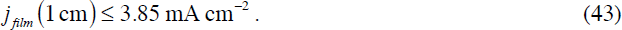

These estimated current densities are within only 1 to 2 orders of magnitude of the estimated maximum potential current density that can be extracted from sunlight (equation 20). If attainable, these could greatly relieve the need for highly extended surface areas inside an electrosynthesis reactor highlighted by the analysis of H_2_-mediated electron transport.

It is worth noting that for the case of 1 cm thick conductive films, the electrode surface area required to host the film is only ≈ 10 times larger than that of the ground upon which the light generating this current is incident. Given that light harvesting in photosynthesis saturates at ≈ 10% of peak solar flux, this area is also approximately equal to that over which crops would need to be spread for the incident light to be reduced to non-saturating levels.

Additionally, it is conceivable that these current densities could be even higher given the potential for higher rates of carbon fixation. For example, elevating the CO_2_ concentration in an electrosynthesis system could lead to substantially higher RuBisCO turnover rates^87,102^. Elevated CO_2_ levels could be further leveraged by utilizing RuBisCO variants with very high turnover rates^103^. Alternatively, the artificial nature of electrosynthesis invites attempts to replace the Calvin cycle with one of the several alternative, potentially faster and more efficient carbon fixation cycles found in nature^104-109^, or computationally designed cycles^110^. This raises the possibility of further compaction of the electrosynthesis process.

Determining if these estimates of current density are achievable, and achievable with low energy losses, will require a deeper physical understanding of the mechanisms of solid matrix EET. These remain contentious, and it is possible that it could vary from species to species. Seminal work by Malvankar *et al.*^111,112^ points to the possibility that *Geobacter* may use pili proteins that act like organic semiconductors or metals. Alternatively, work by Glaven *et al*. in *Geobacter sulfurreducens*^113^ and *Shewanella putrefaciens*^114,115^ and theoretical work by Polizzi *et al*.^116^ indicates that the long range transfer mechanism may be redox gradient driven, with electrons hopping from metal center to metal center. A resolution to this debate would greatly aid attempts to estimate maximum transfer rates and energy losses in solid matrix EET.

An additional challenge to this approach is that many organisms capable of solid matrix conduction including *Geobacter sulfurreducens* and *Clostridium pasteurianum*^117^ are highly O_2_ sensitive. Maintaining an anaerobic environment inside the cathode chamber of the electrosynthesis reactor, while splitting water in the anode chamber and maintaining a low cell resistance remains a formidable engineering challenge, particularly at large scales. This restriction highlights the need for further research into hosts with greater O_2_-tolerance and further investigation into the O_2_-sensitivity of key molecular components of EET.

### 5.2 Transferring Electrons from the Cell Surface to Metabolism In Solid Matrix Conduction

Figure 5B shows a model for electron uptake through a membrane spanning EET pathway. This pathway transfers electrons that were delivered to the cell surface either by direct contact with the electrode or by solid matrix conduction into the inner membrane. The is drawn from data on electron uptake in *Shewanella oneidensis*, but may apply to other systems with membrane spanning EET pathways such as the Mto pathway in *Sideroxydans lithotrophicus* ES-1^47^ and the Pio pathway in *Rhodopseudomonas palustris*^43,51^.

Although *Shewanella* is known primarily for its ability to expel electrons from metabolism Ross *et al.* demonstrated that cathode-donated electrons can traverse the the Mtr complex, enter the quinone pool in the inner membrane and drive periplasmic enzymatic reactions^58^. Similarly, Bose *et al.* demonstrated that the Pio complex, a counterpart to Mtr, is necessary for *R. palustris* to uptake electrons from a cathode.

Although work by Ross *et al*.^58^ demonstrates that electrons injected from an electrode through the Mtr system are capable of entering the quinone pool in the inner membrane, it is unclear, at the time of writing, if these electrons are capable of reducing the NADH and NADPH pools of *S. oneidensis* or of producing a proton gradient needed to regenerate ATP. For *S. oneidensis* the question marks in figure 5B should not be taken just as a question of identity, but one of existence.

The ability of *R. palustris* and *S. lithotrophicus* to fix carbon through the Calvin cycle suggests strongly that in these organisms a mechanism for proton motive force generation and NADH and NADPH pool reduction exists. However, at the time of writing, the exact mechanism remains unknown^118,119^. Understanding this machinery, which one could imagine as being analogous to an electrical power transformer, would make significant contributions to understanding the upper limits of efficiency of electrosynthesis. In addition, knowledge of this machinery could permit organisms such as *E. coli* to acquire electrosynthetic ability. It should be noted, that this “uphill pathway” which raises the energy of electrons and transfers them from the quinone pool (at ≈ -100 mV vs. SHE) to NAD(P)H (at ≈ -320 mV vs. SHE) may represent a large energy loss when compared with the relatively efficient direct transfer of electrons from H_2_ to NADH in H_2_-mediated electron uptake.

## 6. Conclusions

The world is replete with renewable energy and it is conceivable, if not likely, that a significant fraction of world energy use will be sourced from sustainable sources in the coming years. There is enough power floating through air to power the world hundreds of times over, if only it could be captured and stored cheaply.

While the limitations of light capture of natural photosynthesis strongly suggest the efficiency advantages of electrosynthesis, these advantages have yet to be conclusively demonstrated. However, as of the time of writing there is no consensus on the upper limits of efficiency of electrosynthesis. Determining what sets these limits, estimating them, and then building prototype systems that approach these boundaries are key research challenges in electrosynthesis today.

Finally, physics alone cannot prescribe the best course of design for electrosynthesis, but it can help to guide biological and system engineering choices in the construction of microbes and reactors to maximize efficiency and minimize space requirements. The best way to select between these choices and validate the potential of electrosynthesis is to build prototype reactors and organisms and test them in the real world.

## Acknowledgements

This work was supported by a Career Award at the Scientific Interface from the Burroughs Wellcome Fund to Buz Barstow and Princeton University startup funds. The author wishes to thank Professor Andrew Bocarsly for helpful discussions.

### List of Symbols and Abbreviations

**List of Symbols**

*U*: Extractable stored energy per unit area in a given time. Usually expressed in J m^-2^.
d*t*: Infinitesimal time interval.
*t*: Time.
Δ*t*: Finite time interval, usually 1 month or a growing season.
η*o*: Overall energy conversion efficiency. Dimensionless with a value from 0 to 1.
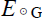: Solar energy delivered during a given time, Δ*t*. Usually expressed in J m^-2^.
*A_T_*: Area required to capture US transportation energy demands from sunlight.
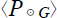: Solar power at ground level, averaged across day and night. Usually expressed in W m^-2^.
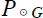: Solar power at ground level. Usually expressed in W m^-2^.
*Y*: Genetic yield. Energy stored under optimal growing conditions in a given time. Usually expressed in J m^-2^.
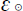: Solar spectrum utilization fraction. Dimensionless. Usually has a value of 0.487.
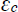: Conversion efficiency. Dimensionless with a value of 0 to 1.
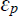: Partitioning efficiency. Dimensionless with a value of 0 to 1.
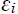: Interception efficiency. Dimensionless with a value of 0 to 1, although with a usual maximum of 0.95.
φ: Latitude.
λ: Longitude.
*U_b_*: Energy content of fixed carbon stored as biomass.
*c_C_*: Carbon content of dry wood. Dimensionless with a value of 0 to 1.
*u_w_*: Energy content of wood. Usually expressed in kilojoules per gram.
m_C_: Net mass of carbon stored by photosynthesis per m_2_ of land during A *t*. Usually expressed in grams of carbon per m_2_.
*L*: Leaf area index. Dimensionless, but expresses in m_2_ of leaf per m_2_ of ground.
*k*: Extinction coefficient of canopy. Dimensionless.
*R_g_*: Radius of gyration. Expressed in m.
*A_i_*: Light interception cross section of cell. Expressed in m_2_.
*N_A_*: Avogadro’s number.
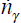: Rate of delivery of photosynthetically active photons to the cell. Expressed in photons per second.
*m_p_*: Dry mass of protein in the cell. Measured in grams.
*m_d_*: Dry mass of cell. Measured in grams.
*f_p_*: Fraction of dry mass constituted by protein
*f_R_*: Fraction of dry protein mass constituted by RuBisCO.
*n_R_*: Number of RuBisCO molecules per cell.
*E_γ_*: Average energy of one mole of PAR photons. Measured in joules.
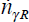: Photon demand rate of carbon fixation in the cell. Measured in photons per second.
*n_cb_*: Number of CO_2_ molecules needed to synthesize one butanol molecule.
*n_eb_*: Number of electrons needed to synthesize one butanol molecule.
*r_R_*: RuBisCO turnover rate. Measured in molecules of CO_2_ per second.
*A_cs_*: Surface area of cylindrical cell. Measured in m_2_.
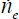: Electron demand rate per cell for butanol synthesis. Measured in electrons per second.
*j*: Current density. Usually measured in μA cm^-2^.
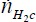: Rate of delivery of photons to the ground with an energy exceeding 1.23 eV, that necessary to split water and reduce protons to H2. Energy difference is that between the redox couples of O_2_/H_2_O (0.82 V vs. SHE) and H^+^/H_2_ (-0.41 eV vs. SHE).
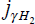: Current density corresponding to 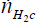.
*r_H_*: Hydrogen oxidation turnover rate of hydrogenase enzyme.
*n_H_*: Number of hydrogenases needed to satisfy electron demands of carbon fixation.
*M_H_*: Total mass of hydrogenases in cell.
*ϱ_c_*: Cell density.
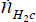: Total H_2_ demand rate for a culture of cells.
*j*_H_2__: Effective current density for H_2_-mediated electron transport.
*j_c_*: Current density demand of single cell in solid matrix conduction.

**Abbreviations**

EJ: Exajoules or 10_18_ joules. Equivalent to 1.055 quadrillion British Thermal Units (quads).
TW: Terawatts or 10_12_ watts.
EET: Extracellular electron transport.
PAR: Photosynthetically Active Radiation. Portion of the solar spectrum from 400 to 740 nm.
SHE: Standard Hydrogen Electrode. Electrochemical reference energy level.

